# The haploinsufficient tumor suppressor Tip60 negatively regulates the oncogenic Aurora B kinase

**DOI:** 10.1101/589473

**Authors:** Arnab Bose, Surabhi Sudevan, Vinay J. Rao, Hiroki Shima, Kazuhiko Igarashi, Tapas K. Kundu

## Abstract

The Aurora kinases represent a group of serine/threonine kinases which are crucial regulators of mitosis. Dysregulated Aurora kinase B (AurkB) expression, stemming from genomic amplification, increased gene transcription or overexpression of its allosteric activators, is capable of initiating and sustaining malignant phenotypes. Although AurkB level in cells is well-orchestrated, studies that relate to its stability or activity, independent of mitosis, are lacking. We report that AurkB undergoes acetylation *in vitro* by lysine acetyltransferases (KATs) belonging to different families, namely by p300 and Tip60. The haploinsufficient tumor suppressor Tip60 acetylates two highly conserved lysine residues within the kinase domain of AurkB which not only impinges the protein stability but also its kinase activity. These results signify a probable outcome on the increase in “overall activity” of AurkB upon Tip60 downregulation, as observed under cancerous conditions. The present work, therefore, uncovers an important functional interplay between AurkB and Tip60, frailty of which may be an initial event in carcinogenesis.

## 1. Introduction

AurkB is one of the three human paralogs of Aurora kinases and is a member of the chromosome passenger complex. It plays pivotal roles in chromosome congression, spindle bi-orientation and cytokinesis (Ruchaud S *et al.* 2007) and undergoes a multitude of post translational modifications which modulate its spatio-temporal behaviour necessary for proper mitotic progression (Monaco L *et al.* 2005; Ban R *et al.* 2011; Fernández-Miranda G *et al.* 2010; Catherine Lindon *et al.* 2016). Although *AurkB* transcription or protein stability is tightly regulated prior to mitotic entry or following mitotic exit respectively (Kimura M *et al.* 2004; Park J *et al.* 2018), this regulation is lost in cancer cells, leading to elevated steady state AurkB protein levels. It is frequently overexpressed in both solid and haematological malignancies, and is therefore a potential target of ani-neoplastic therapies (Borisa AC *et al.* 2017).

Proteome-wide high throughput studies have unveiled the “lysine acetylome” of a cell (Choudhary C *et al.* 2009). Interestingly, many cell cycle regulators were found to be acetylated. Congruently, histone deacetylase (HDAC) inhibitors were found to affect kinetochore assembly through decreased pericentromeric targeting of AurkB (Robbins AR *et al.* 2005). HDAC3 was observed to not only assist in histone H3 deacetylation during mitosis to provide a hypoacetylated H3-tail for phosphorylation by AurkB (Li Y *et al.* 2006) but also deacetylate AurkB and enhance its kinase activity (Fadri-Moskwik M *et al.* 2012). Additionally, AurkB interacts with class IIa histone deacetylases in determining their nuclear localization during mitosis (Guise AJ *et al.* 2012).

We observe that p300 and Tip60 acetylate AurkB *in vitro*. Tip60 mediated acetylation of AurkB at two highly conserved K85 and K87 residues negatively influence its protein stability and kinase activity. We propose that under normal physiology Tip60 mediated acetylation serves to contain AurkB oncogenicity. However, upon Tip60 downregulation, as observed in cancers (Gorrini C et al. 2007; Mattera L *et al.* 2009), AurkB is not only stabilized but also exhibit increase in its kinase activity, causing an overall increase in activity of the oncogenic kinase.

## 2. Materials and Methods

### 2.1. Cell culture, antibodies and reagents

HEK293 and HEK293T (purchased from ATCC), MDA-MB-231 and MCF7 were grown at 37°C in DMEM (Invitrogen, Thermo Fisher Scientific) containing 10% fetal bovine serum. To generate Flag-AurkB overexpressing stable cell line, pCDH adenoviral vector containing N-terminal Flag-AurkB fusion was co-transfected with lentiviral gene containing plasmids (psPAX2, pRSV-Rev and pCMV-VSV-G) into HEK293T cells to package the mammalian clone into the viral particles. Forty-eight hours post-transfection, the generated viral particles were used to infect HEK293 cells, and subsequently sorted for the GFP-positive cell population. Constitutive Tip60 knockdown cell line and the corresponding control cell line were similarly established in HEK293 using Tip60 directed shRNA or a non-silencing shRNA cloned in pGIPZ vector (Dharmacon). Transient transfection of 2N3T-Tip60 or the corresponding control empty vector was carried out for 48 hours into HEK293 cells using Lipofectamine 2000 transfection reagent (Invitrogen); UV irradiation was carried out at a dose of 15mJ/cm^2^ using Stratagene Stratalinker 1800; HEK293 cells overexpressing Flag-AurkB was treated with 10 µM of trichostatin A (TSA) (Sigma-Aldrich) or DMSO for 12 hours. Cells for the various treatments or transfection conditions were harvested by scraping and lysed in lysis buffer containing 50mM tris-HCl, pH 7.4, 150mM NaCl, 5mM EDTA, 1% NP40, and 1 X protease inhibitory cocktail (Sigma-Aldrich). Anti-Aurora kinase B antibody was purchased from Abcam; anti-Flag, anti-β-actin and anti-HA antibodies were purchased from Sigma-Aldrich; anti-tubulin antibody was purchased from Millipore.

### 2.2. Cycloheximide chase assay

3 X Flag pCMV10-AurkB WT or the corresponding acetylation defective (K85R-K87R) and mimetic (K85Q-K87Q) mutants were transiently transfected into HEK293 cells in 60mm dish format using Lipofectamine 2000. Twenty-four hours post transfection, the cells were trypsinized and re-seeded into 6 well-plate and allowed to grow for 24-36 hours more to grow till 60-70% confluence, after which they were treated with 150 µg/ml of cycloheximide (Sigma-Aldrich) for the indicated time periods. The cells were harvested by scraping and lysed in Laemmli buffer.

### 2.3 *In vitro* lysine acetyltransferase (KAT) assay

KAT assay was performed using highly purified, baculovirus-expressed, recombinant full-length lysine acetyltransferases-His_6_-Tip60, Flag-PCAF or His_6_-p300 from Sf21 insect cells as enzymes and His_6_-AurkB, as substrates, in a 30 μl final reaction volume. Briefly, the reaction mixture consisting of 50 mM tris-HCl, pH 8.0, 10% (v/v) glycerol, 1 mM dithiothreitol (DTT), 1 mM phenyl methyl sulfonyl fluoride (PMSF), 0.1 mM EDTA, pH 8.0, 10 mM sodium butyrate, and 1 μl of 3.3 Ci/mmol of H^3^-acetyl Coenzyme A (acetyl-CoA) and the indicated proteins, was incubated at 30°C for 30 min. Mass acetylation was carried for 2.5 hours at 30°C (with replenishment of indicated enzymes and acetyl-CoA every 30 mins, thrice). To visualize the radiolabelled acetylated protein, the reaction products were resolved on 12% SDS-polyacrylamide gel upon electrophoresis and stained by coomassie brilliant blue (CBB) to ascertain the presence of proteins. The gel was then dehydrated in DMSO for 1hr and incubated with scintillation fluid (22.5% w/v PPO solution in DMSO) for 30 minutes and finally equilibrated in water for two hours. The rehydrated gel was dried using a gel drier and exposed to X-ray film for 10-12 days.

### 2.4. *In vitro* kinase assay

Recombinant histone H3, expressed and purified from *E. coli*, was incubated with either mass acetylated (Tip60-mediated) or mock acetylated His_6_-Aurora kinase B, in a 30 µl reaction mixture containing 50 mM tris-HCl, 100 mM NaCl, 0.1 mM EGTA, 10 mM MgCl2, 0.2% β-mercaptoethanol and [γP32] ATP (Specific Activity 3.5 Ci/mmol). The reaction mixture was incubated at 30°C for 15 minutes. The reaction was inhibited over ice for 5 minutes, constituent proteins were denatured by the addition of gel loading dye containing SDS, heated at 90°C for 5 minutes and resolved using 12% denaturing PAGE, stained by CBB, and followed by autoradiography using X-ray films.

## 3. Results and Discussions

### 3.1. Tip60 acetylates AurkB at two highly conserved lysine residues

In order to identify the putative KAT(s) responsible for the lysine acetyltransferase activity toward AurkB, we adopted an unbiased approach and considered the screening of KATs belonging to three different families of lysine acetyltransferases, namely p300, PCAF and Tip60. Each of these enzymes were purified from Sf21 insect ovarian cells using suitable baculoviral constructs containing the human KAT homolog. Recombinant His_6_-AurkB, expressed and purified from *E. coli* was used for an *in vitro* lysine acetyltransferase (KAT) assay. It was intriguing to find that p300 and Tip60 were capable of acetylating AurkB, while PCAF could not (figure 1).

**Figure 1.**
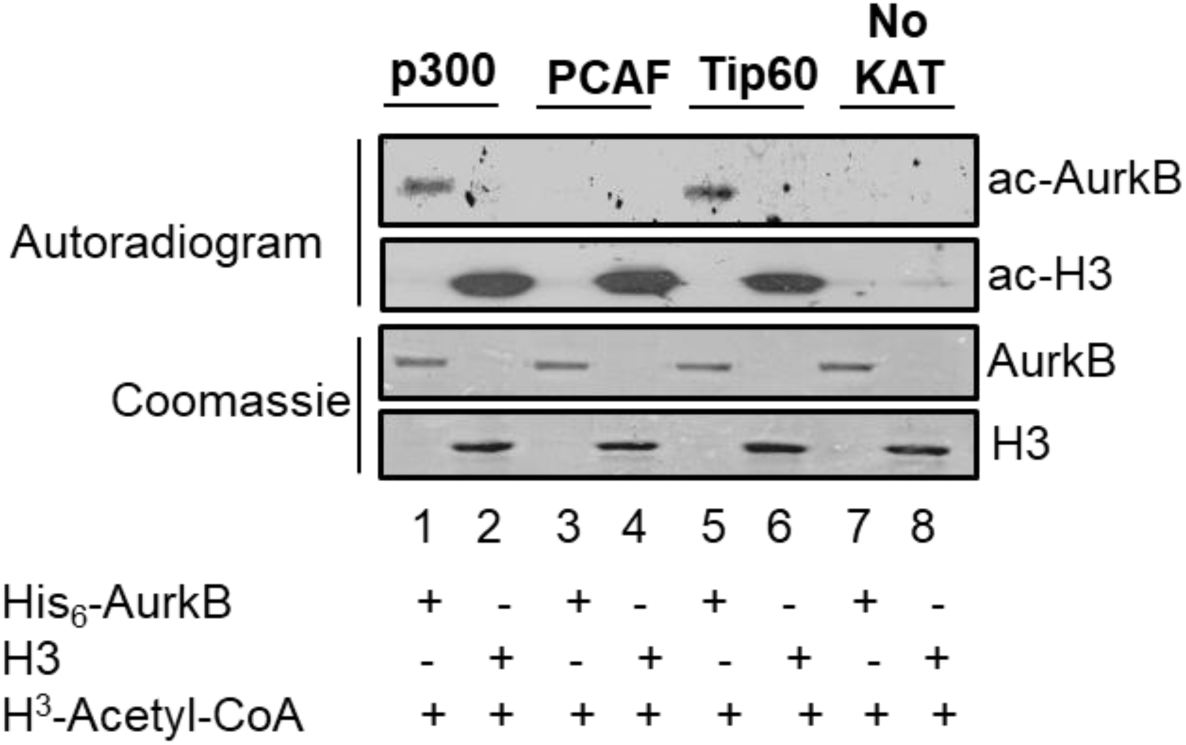
Tip60 and p300 acetylate AurkB. 500ng each of recombinant His_6_-AurkB and histone H3, purified from *E. coli*, were subjected to *in vitro* lysine acetyltransferase (KAT) assay by using His_6_-p300, Flag-PCAF and His_6_-Tip60 purified from Sf21 cells using suitable baculoviral constructs and tritiated acetyl-CoA (H^3^-acetyl-CoA). No KAT (lanes 7 and 8) served as the negative control in which none of the KATs were added.

The haploinsufficient tumor suppressor, Tip60 is known to oppose oncogenesis. Factual downregulation of Tip60 expression in multiple cancer types and its direct involvement in acetylation dependent inhibition of oncogenic pathways (Du Z *et al.* 2010; Kim MY *et al.* 2007) led us to study the further possibility of a Tip60 dependent regulation of AurkB in the context of acetylation. In order to identify the acetylation sites, we performed *in vitro* mass acetylation of recombinant full length His_6_-AurkB using His_6_-Tip60. The acetylated protein was subjected to liquid chromatography-coupled tandem mass spectrometry (LC-MS/MS) and nine probable acetylated lysine residues were identified-K4, K31, K56, K85 (supplementary figure 1, panel-I), K87 (supplementary figure 1, panel-II), K115, K168, K195, and K202. Additionally, we observed that K85 and K87 residues reside in a glycine-rich microenvironment and is part of a G-K-G-K motif, which is similar to G-K-X-G-K motif found in the well-studied Tip60 substrates-histone H4 and Twist (figure 2A). Noting the high extent of conservation of these two lysine residues (figure 2B), we mutated them, either singly or in combination, to arginine and the *E. coli* purified recombinant WT and mutant AurkB proteins were subjected to *in vitro* KAT assay. While the K85R-K87R double point mutant led to a complete abrogation of Tip60 mediated acetylation, the single point mutants were only minimally affected (figure 2C). The K85R-K87R mutant did not show any acetylation even for higher Tip60 concentrations (figure 2D), emphasizing these sites to be the major Tip60 mediated acetylation sites.

**Figure 2.**
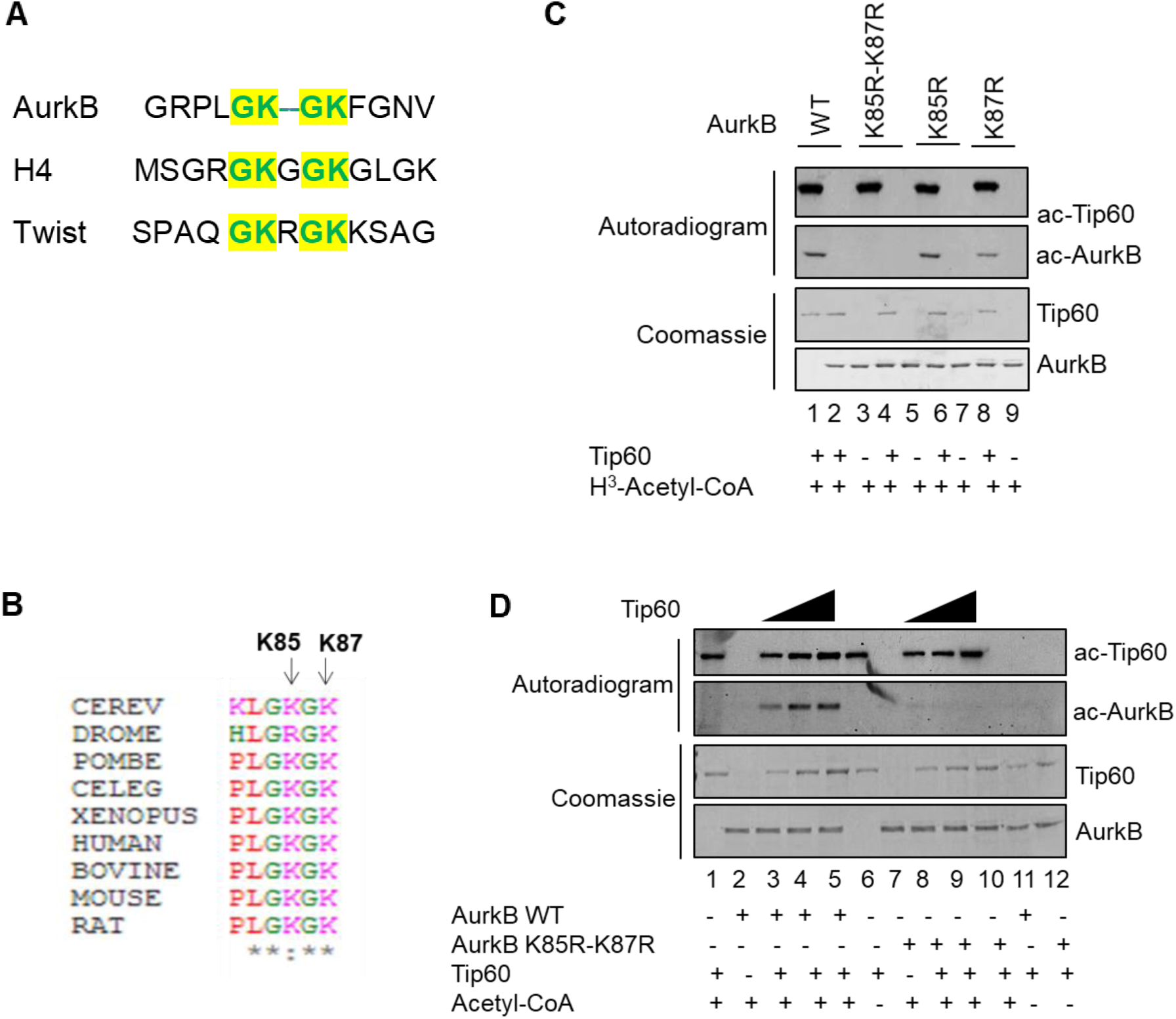
Tip60 acetylates AurkB at K85 and K87. **A.** Local alignment of oligopeptide sequence displaying the similarities of putative Tip60 mediated acetylation motif on AurkB, histone H4 and Twist. **B.** Local alignment exhibiting the extent of conservation of K85 and K87 residues of AurkB in different organisms-CEREV, Saccharomyces cerevisiae; DROME, Drosophila melanogaster; POMBE, Schizosaccharomyces pombe; CELEG-Caenorhabditis elegans; XENOPUS, Xenopus laevis; HUMAN, Homo sapiens; Bovine, Bos Taurus; Mouse, Mus musculus; RAT, Rattus rattus. **C.** 500ng each of His_6_-AurkB WT or the indicated mutant His_6_-AurkB proteins were used to carry out an *in vitro* KAT assay using His_6_-Tip60 purified from Sf21 cells and H^3^-acetyl coenzyme A. **D.** 500ng each of His_6_-AurkB WT or His_6_-AurkB K85R-K87R double point mutant proteins were used to carry out an in vitro KAT assay using increasing concentrations of His_6_-Tip60 purified from Sf21 cells and H^3^-acetyl coenzyme A. The coomassie staining for each of the lanes is shown below the autoradiogram profiles for comparing the loading levels across each lane.

### 3.2. Tip60 and p300 may not possess overlapping acetylation sites on AurkB

The screening of different KATs had highlighted that p300, apart from Tip60, is also capable of acetylating AurkB *in vitro*. We subsequently studied the acetylation of K85 and K87 residues of AurkB by p300 and observed that p300 is capable of acetylating either of the individual point mutants-K85R and K87R (figure 3A) or the K85R-K87R double mutant (figure 3B). These results emphasize either the exclusivity of K85 and K87 residues toward Tip60 mediated acetylation or portrays the possibility of numerous p300 mediated acetylation sites on AurkB, and mutation of two (K85 and K87) amongst many, is incapable of altering the overall p300 dependent acetylation status of the AurkB protein.

**Figure 3.**
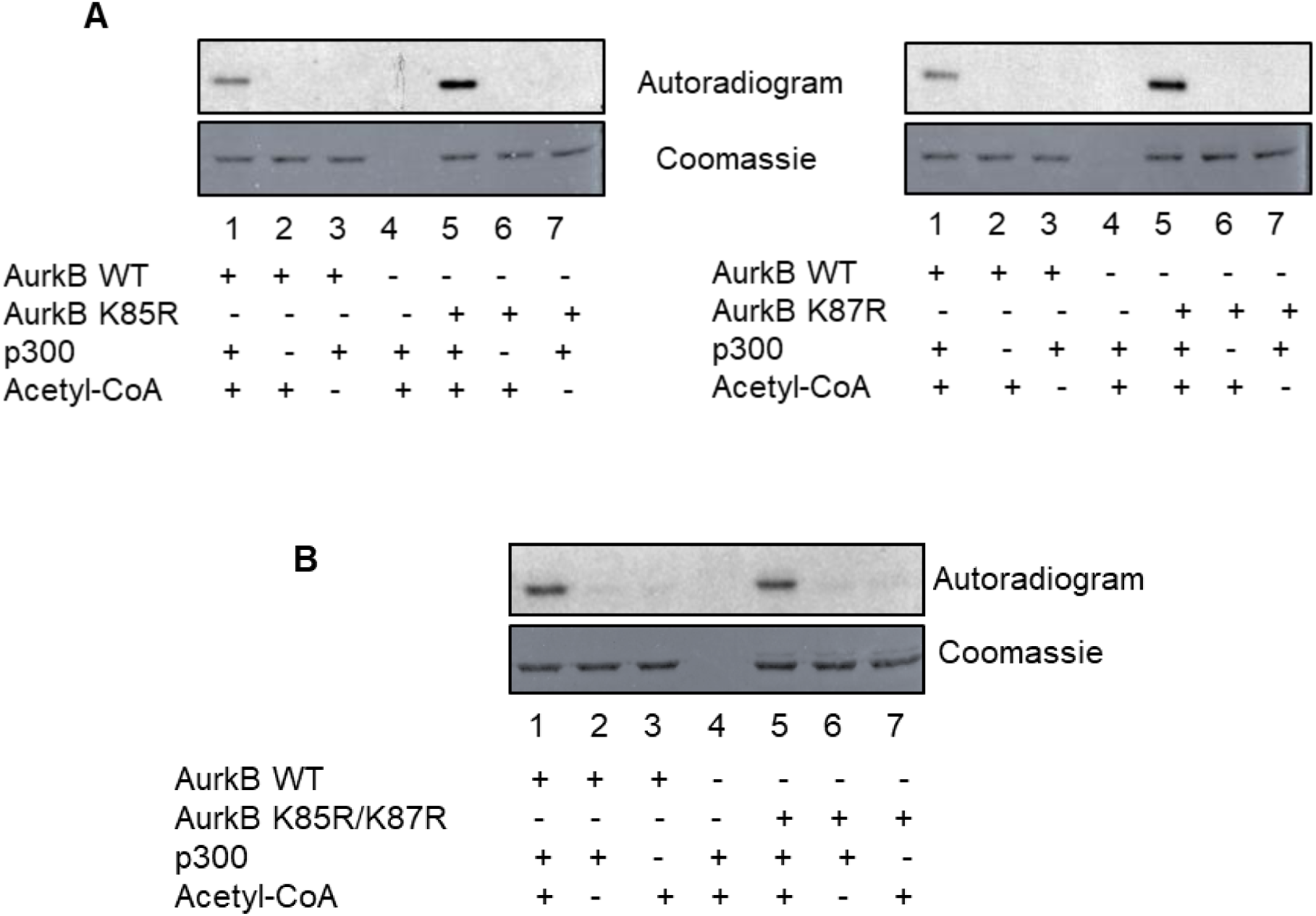
Validation of p300 mediated acetylation of AurkB for Tip60 targeted lysine residues. 500ng each of recombinant His_6_-AurkB WT or the His_6_-AurkB mutant proteins (single point mutants-K85R and K87R, as in panel **A**) or the double point mutant (His_6_-AurkB K85R/ K87R, as in panel **B**) were used to carry out an *in vitro* KAT assay using His_6_-Tip60 purified from Sf21 cells and H^3^-acetyl coenzyme A. The coomassie staining for each of the lanes is shown below the autoradiogram profiles for comparing the loading levels across each lane.

### 3.3. Tip60 interacts with AurkB in cells

We studied if AurkB and Tip60 interact with each other in a cellular context. Reciprocal co-immunoprecipitation of ectopically overexpressed 2N3T-Tip60 (henceforth referred to as HA-Tip60) in HEK293 cells constitutively expressing Flag-AurkB confirmed their physical interaction (figure 4). These results demonstrate that Tip60 interacts with AurkB in cells and acetylate it at K85 and K87 residues.

**Figure 4.**
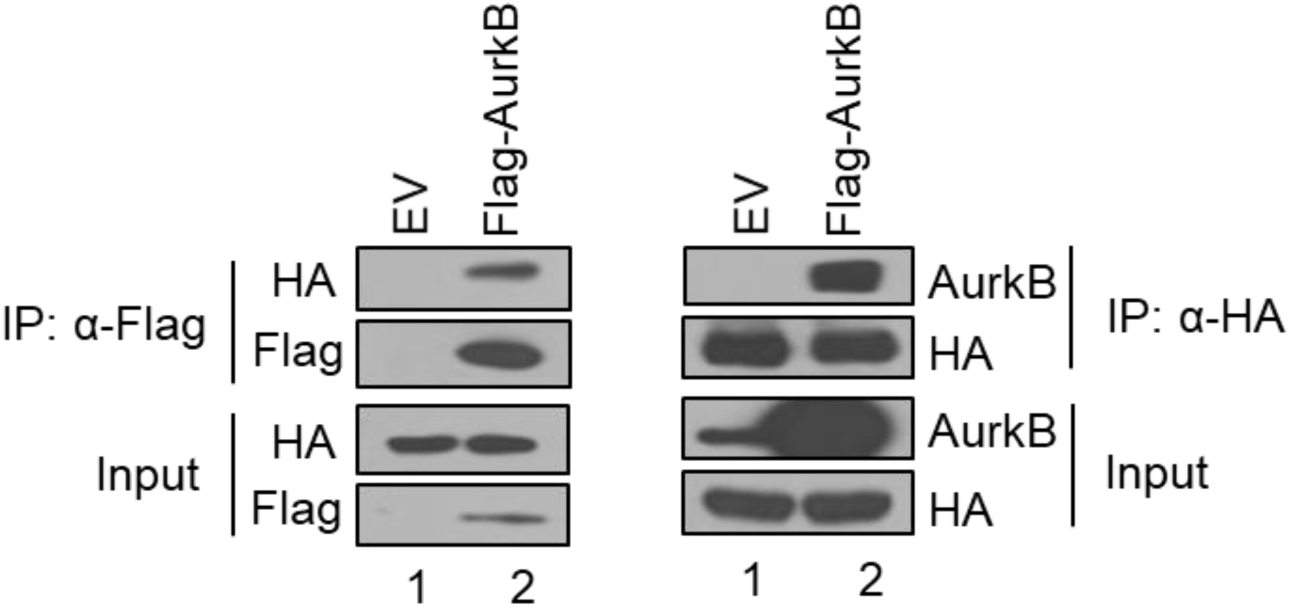
Tip60 interacts with and acetylates AurkB in cells. Reciprocal co-immunoprecipitation showing the interaction between HA-Tip60 (transiently overexpressed for 48 hours) and Flag-AurkB in HEK293 cells constitutively expressing Flag-AurkB protein.

### 3.4. Tip60 regulate protein levels of AurkB in cells

Histone deacetylases play important roles in mitosis. Broad and non-selective HDAC inhibitors like trichostatin A (TSA) inhibits class I HDACs (HDAC1, 3, 4, 6, and 10) and has been shown to induce G2/M arrest by targeting HDAC3 (Yun Li *et al.* 2006). Moreover, HDAC3 was also reported to deacetylate AurkB (Fadri-Moskwik M *et al.* 2012). We wanted to study the effect of TSA treatment on the levels of AurkB. Strikingly, treatment of HEK293 cells stably expressing Flag-AurkB with TSA led to a marked reduction of the AurkB protein pool (figure 5A). We wondered about the likelihood of acetylation of AurkB as a trigger for its downregulation. To this end, we overexpressed HA-Tip60 in HEK293 cells and found that the endogenous AurkB levels exhibited a marked reduction, which could be rescued by simultaneous Tip60 knockdown (figure 5B, panel-I; compare lane 1 v/s 2 and 2 v/s 3). Similar rescue was also observed when Flag-AurkB was transiently overexpressed in constitutive Tip60 knockdown cells (supplementary figure 2; figure 5B, panel-II). We could faithfully reproduce these results in two other breast cancer cell lines, MDA-MB-231 and MCF7 (figure 5C, panel-I and II, respectively]. We also generated point mutants corresponding to acetylation defective (K85R-K87R) or mimetic (K85Q-K87Q) conditions, on Flag-AurkB mammalian expression vector, and observed that the later exhibited reduced stability under cycloheximide chase assay conditions (figure 5D). These observations collectively confirm that Tip60 mediated acetylation of AurkB at K85 and K87 residues result into its destabilization in cells.

**Figure 5.**
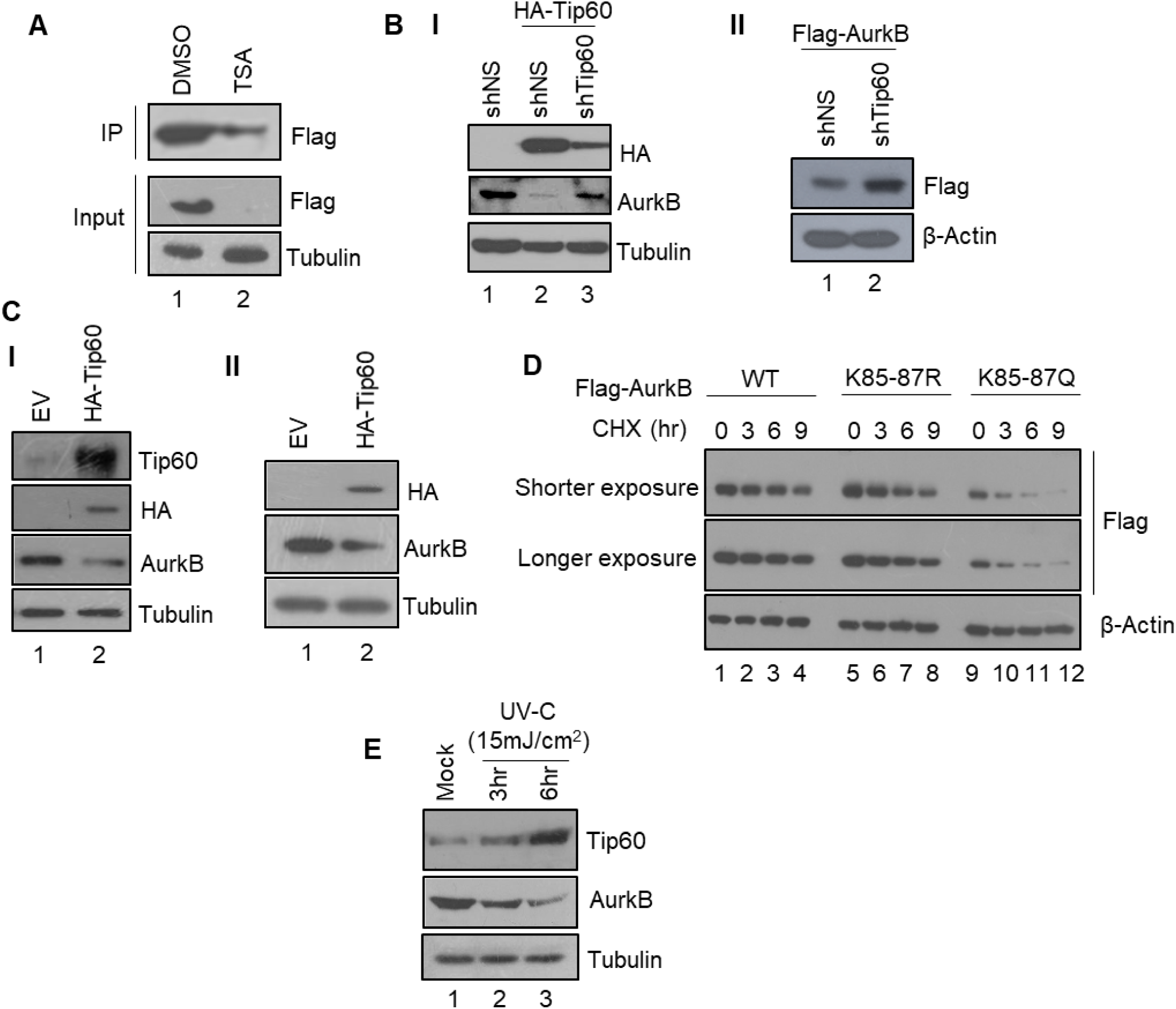
Tip60 regulates AurkB levels in cells. **A.** Western blot representation of the indicated proteins upon treatment of stably overexpressing Flag-AurkB HEK293 cells with TSA or the corresponding DMSO control. IP represent the immunoprecipitated fraction from 1mg of whole cell lysate; input represent the whole cell lysate fraction. **B.** Western blot analysis showing that Tip60 overexpression reduces, while simultaneous Tip60 knockdown rescues AurkB levels in HEK293 cells **(panel-I)** [shNS represent non-silencing shRNA control]. **(Panel-II)** Transient overexpression of Flag-AurkB in stable shNS and shTip60 cell lines showing the higher levels of Flag-AurkB in the Tip60 knockdown cell line as compared to the control shNS cells. **C.** Western blot representation for overexpression of Tip60 in MDA-MB-231 cells **(panel-I)** or MCF7 cells **(panel-II)** and study of the proteins with the indicated antibodies. **D.** Cycloheximide chase assay for WT, acetylation defective (K85R-K87R) or acetylation mimetic (K85Q-K87Q) mutants of AurkB. **E**. Western blot representation of HEK293 cells exposed to 15mJ/cm^2^ UV-C (254 nm) and harvested at the indicated time points after irradiation for the indicated proteins.

We next asked if AurkB modulation agrees with a physiologically relevant scenario where Tip60 exhibits differential expression levels in cells. An earlier observation had demonstrated enhanced stabilization of Tip60 protein in cells upon UV exposure (Legube G *et al.* 2002), owing to an inhibition of its degradation by MDM2. In agreement with this study, we observed that UV irradiation caused enhancement in Tip60 levels in cells and AurkB was concomitantly downregulated under such conditions (figure 5E).

### 3.5. Tip60 mediated acetylation inhibit the kinase activity of AurkB

K85 and K87 residues lie within the kinase domain of AurkB. This propelled us to study if the kinase activity is affected upon acetylation by Tip60. Recombinant His_6_-AurkB was acetylated in vitro with Tip60 and the acetylated kinase was used in an in vitro kinase assay with histone H3 as a substrate. We found that acetylation of AurkB lowered its kinase activity, when compared to the mock acetylated control (supplementary figure 3; figure 6A). We extended our study in assessing the point mutants of AurkB and observed that the acetylation mimetic K85Q-K87Q mutant exhibited reduced kinase activity as compared to either the WT or the acetylation defective K85R-K87R mutant (figure 6B). The acetylation defective mutant, however, showed higher kinase activity, even as compared to the WT kinase, the cause of which is presently unclear. These results confirm that Tip60 dependent acetylation inhibit the kinase activity of AurkB.

**Figure 6.**
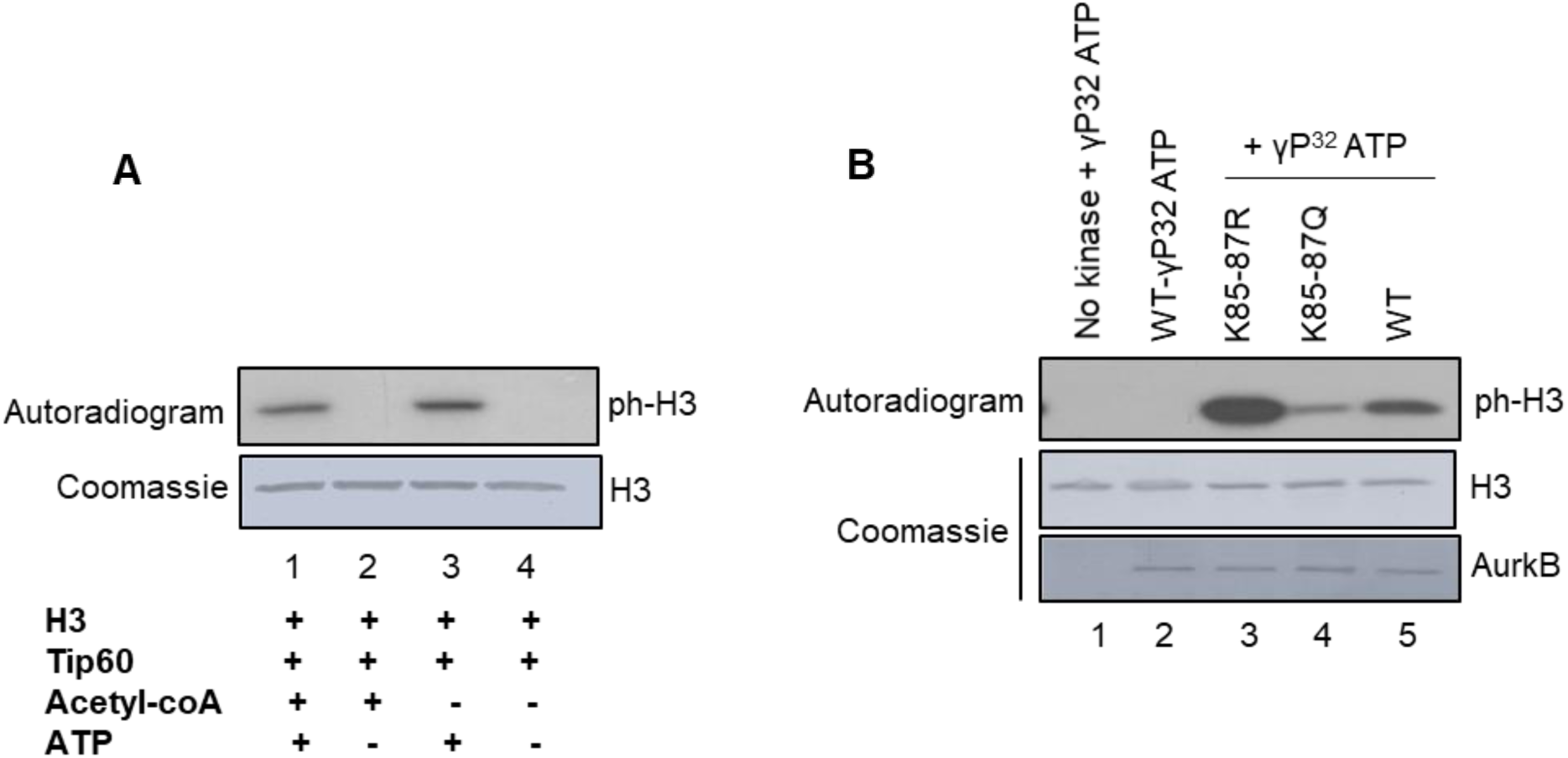
Tip60 mediated acetylation inhibits AurkB activity. **A.** 500ng of His_6_-AurkB was either acetylated with Tip60 or mock acetylated (control) in vitro. One-tenth of the acetylation reaction was used to carry out in vitro kinase assay with 500 ng of recombinant H3. The coomassie staining for each of the lanes is shown below the autoradiogram profiles for comparing the loading levels across each lane. **B.** *In vitro* kinase assay was carried out using 100ng each of WT, acetylation defective mutant (K85R-K87R), or acetylation mimetic mutant (K85Q-K87Q) of AurkB using histone H3 as a substrate. Lesser extent of kinase activity was observed for the K85Q-K87Q mutant as compared to either the WT or the K85R-K87R mutant.

## 4. Discussion

AurkB is the catalytic component of the chromosomal passenger complex (CPC) and ensures proper chromosome alignment and segregation (Adams RR *et al*. 2001). It also assists in cleavage furrow formation, and cytokinesis and hence a reduction in the kinase activity by either chemical inhibitors or RNA interference gives rise to chromosome segregation defects and aneuploidy (Ditchfield C *et al.* 2003; Hauf S *et al.* 2003). AurkB overexpression in cultured mammalian cells and mouse model is reported to cause an increased tumor incidence by tetraploidization or inhibiting p53 and p21 levels and functions (Gully CP *et al.* 2012; González-Loyola A *et al.* 2015; Nguyen HG *et al.* 2009) or by modulation of Myc dependent tumors (den Hollander J *et al.* 2010; Yang D *et al.* 2010). Furthermore, the paralogue-AurkA, has been reported to participate in varying signalling events which culminate into tumorigenesis and cancer progression (Katayama H *et al.* 2004; Briassouli P *et al.* 2007; Dar AA *et al.* 2009; Otto T *et al.* 2009).

Here, we report a previously unanticipated link between AurkB and Tip60 whereby Tip60 contributes to its tumor suppressive effects by not only inhibiting the kinase activity of AurkB but also through destabilization of the AurkB protein, through acetylation of two highly conserved lysine residues present in the kinase domain of AurkB. We propose that widespread Tip60 downregulation in cancers favour the deregulated increase in the overall activity of AurkB in tumor cells that may act as a causative factor for the associated alterations of p53 or c-Myc pathways. Furthermore, as earlier studies elucidate the direct links between Tip60 and genomic aberrations (Bassi C *et al.* 2016), it is worthwhile to study if any of such effects are mediated by the altered activities of AurkB.

## Acknowledgement

The authors thank Prof. Didier Trouche (LBCMCP, Centre de Biologie Intégrative (CBI), Université de Toulouse, CNRS, UPS, Toulouse, France) for 2N3T empty vector and 2N3T-Tip60 plasmids and Dr. Amit Dutt (Integrated Cancer Genomics Lab, ACTREC, Mumbai, India) for sharing MDA-MB-231 and MCF7 breast cancer cells.

## Abbreviations

AurkB: Aurora kinase B;
Tip60: HIV-1 Tat Interacting Protein, 60kDa;
KAT: lysine acetyltransferase

---

**Supplementary figure 1.**
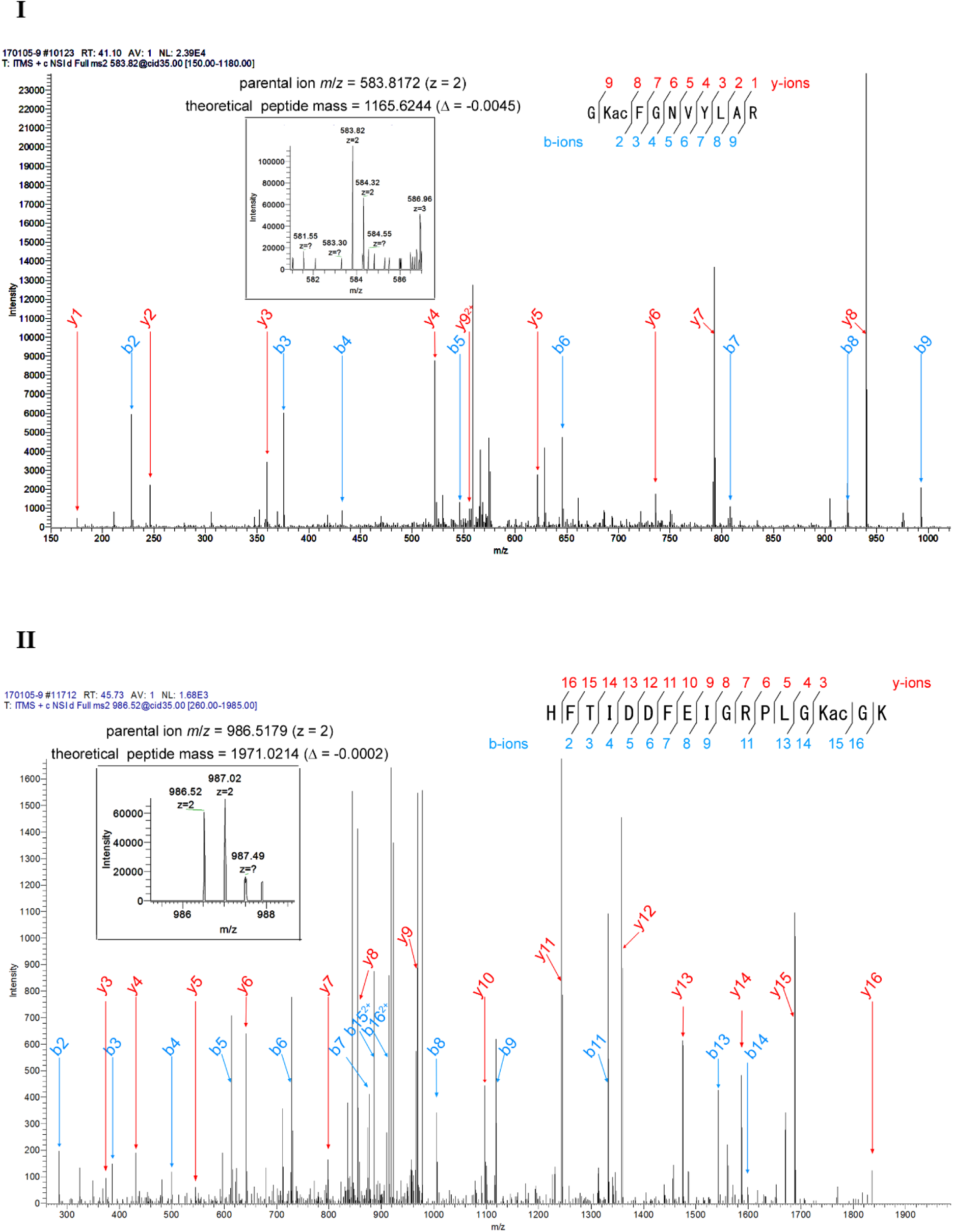
Mass spectra of K85 (panel-I) and K87 acetylation (panel-II).

**Supplementary figure 2.**
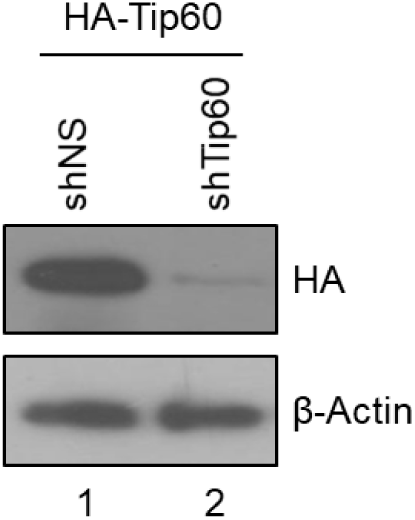
Characterization of constitutive knockdown of Tip60 in HEK293 cells. 2µg of pCMV 2N3T-Tip60 was transiently transfected into either shNS (non-silencing control) or shTip60-HEK293 cells for 48 hours and probed for the indicated proteins.

**Supplementary figure 3.**
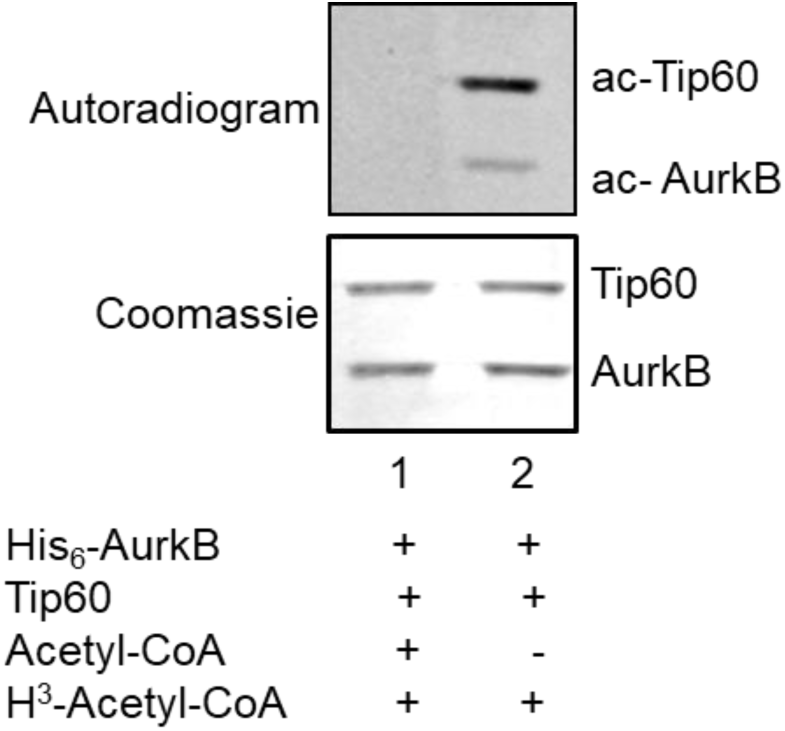
Confirmation of *in vitro* mass acetylation of His_6_-AukB by Tip60. 500ng of His_6_-AurkB was subjected to in vitro mass acetylation by Tip60. After completion of the reaction, 1/10^th^ of the reaction fraction was used for carrying out *in vitro* kinase assay of histone H3; H ^3^-acetyl-CoA was added to the rest of the reaction mixture and the reaction carried out for 30 mins at 30°C. Autoradiogram representation confirms the completion of mass acetylation in lane 1.

